# Brain-inspired methods for achieving robust computation in heterogeneous mixed-signal neuromorphic processing systems

**DOI:** 10.1101/2022.10.26.513846

**Authors:** Dmitrii Zendrikov, Sergio Solinas, Giacomo Indiveri

## Abstract

Neuromorphic processing systems implementing spiking neural networks with mixed signal analog/digital electronic circuits and/or memristive devices represent a promising technology for edge computing applications that require low power, low latency, and that cannot connect to the cloud for off-line processing, either due to lack of connectivity or for privacy concerns. However these circuits are typically noisy and imprecise, because they are affected by device to device variability, and operate with extremely small currents. So achieving reliable computation and high accuracy following this approach is still an open challenge that has hampered progress on one hand and limited widespread adoption of this technology on the other. By construction, these hardware processing systems have many constraints that are biologically plausible, such as heterogeneity and non-negativity of parameters. More and more evidence is showing that applying such constraints to artificial neural networks, including those used in artificial intelligence, promotes robustness in learning and improves their reliability. Here we delve even more in neuroscience and present network-level brain-inspired strategies that further improve reliability and robustness in these neuromorphic systems: we quantify, with chip measurements, to what extent population averaging is effective in reducing variability in neural responses, we demonstrate experimentally how the neural coding strategies of cortical models allow silicon neurons to produce reliable signal representations, and show how to robustly implement essential computational primitives, such as selective amplification, signal restoration, working memory, and relational networks, exploiting such strategies. We argue that these strategies can be instrumental for guiding the design of robust and reliable ultra-low power electronic neural processing systems implemented using noisy and imprecise computing substrates such as subthreshold neuromorphic circuits and emerging memory technologies.

## 1 Introduction

With the advent of deep networks for artificial intelligence [1], and the increasing need of special purpose low-power devices that can complement general-purpose power-hungry computers in “edge computing” applications [2, 3], several types of event-based approaches for implementing Spiking Neural Networks (SNNs) in dedicated hardware have been proposed [4–9]. While many of these approaches are focusing on supporting the simulation of large scale SNNs [4, 6, 9], on converting rate-based Artificial Neural Networks (ANNs) into their spike-based equivalent networks [10–12], or on processing digitally stored data with digital hardware implementations [13–17], the original *neuromorphic engineering* approach, first introduced in the early’90s, proposed to implement biologically plausible SNNs by exploiting the physics of subthreshold analog Complementary Metal-Oxide-Semiconductor (CMOS) circuits to directly emulate the bio-physics of biological neurons and synapses [18, 19].

Today this approach has been extended to include emerging nano-scale memory technologies and a wide range of different types of memristive devices [20–25]. While more difficult to control, due to the analog and noisy nature of the subthreshold analog circuits, and the variability of the memristive devices, this approach has the potential of leading to the construction of extremely compact and low-power brain-inspired neural processing systems [26–29]. Neuromorphic processors built following this approach typically comprise many spiking neuron and dynamic synapse circuits that can be configured to carry out complex spike-based signal processing and learning tasks in real-time. Similar to the biological neural systems they model, these types of neuromorphic systems are extremely low power, operate in a massively parallel fashion, and process information using both analog and asynchronous digital processing methods [28, 30]; they are adaptive, fault-tolerant, and can be configured to display complex behaviors by combining multiple instances of neural computational primitives. On the other hand, as these types of systems are radically different from standard computing platforms based on the Turing machine concept and the von Neumann architecture, there is no well established formalism for “programming them” to carry out pre-defined procedures. Furthermore, due to their analog, continuous-time, and in-memory computing nature, they do not use bit-precise Boolean logic operations, they cannot represent signals with arbitrary precision, and do not support the storage of large amounts of state variables in dense and compact Dynamic Random Access Memory (DRAM) memory blocks. In particular, like their biological counterparts, their processing elements, such as the neuron and synapse analog circuits, are strongly affected by variability and device mismatch [31–33]. All these properties pose significant challenges for understanding how to use this technology to carry out robust computation, in face of the device variability, and without being able to use the classical formalism of computation based on Turing machines.

In this paper we address the problem of achieving robust and reliable computation using an underlying hardware that is noisy and highly variable. This work is in-line with the recent investigations that analyze the role of variability and heterogeneity in neural computation and that attempt to exploit it to for improving learning performance [34–36]. Rather than attempting to minimize the effects of device variability with brute force approaches, we propose brain-inspired computational strategies to counteract the detrimental effects of heterogeneity on spike based computation. We present neural computing primitives that use these strategies for representing and processing signals in a robust manner, and validate them using CMOS neuromorphic chips designed following the original neuromorphic engineering approach [18]. In particular we demonstrate how the use of population coding [37], Excitatory-Inhibitory (E-I) balanced networks [38–40], and Winner-Take-All (WTA) architectures [41, 42] can be exploited for controlling the precision of the signals in these neuromorphic circuits, and we demonstrate how the choice of using signal representations based on population codes, and brain-inspired computational primitives leads to important additional advantages in terms of speed of computation, coding efficiency, and power consumption.

In the next Section we describe the types of neuromorphic systems that we use in this study and quantify the amount of variability present in the neurons, due to device mismatch. In Section 3 we demonstrate how brain-inspired strategies can effectively reduce variability effects, with quantitative measurements, as a function of population size and integration time. Furthermore we show how such strategies offer additional computational advantages, for example in increasing the precision of variable representations, in restoring signals, or in implementing working memory and state-dependent computation. In Section 4 we discuss the benefits of adopting the principles of neural design [43] that we demonstrated with experimental data, and in Section 5 we present the concluding remarks.

## 2 Heterogeneous neuromorphic processing systems

Heterogeneous neuromorphic processing systems typically comprise analog circuits that carry out neural computation, and digital circuits that convert the neuron spikes into digital pulses and transmit them using asynchronous communication schemes. Typically, these systems have been designed using standard CMOS technologies, but with the advent of memristive devices [44, 45], hybrid CMOS-memristive neuromorphic processors started to be proposed as well [21, 23, 46–48]. Examples of pure CMOS mixed-signal neuromorphic systems include the Learning Attractor Neural Network with plasticity and long-term memory chip (LANN-21) [49], the “Final Learning Attractor Neural Network” (F-LANN) chip [50], the Learning-Enabled Neuron Array IC (LENA-IC) [51], the Integrate-and-Fire Array Transceiver (IFAT) architecture [52], the NeuroGrid system [53], the Recurrent On-Line Learning Spiking (ROLLS) neuromorphic processor [54], the Dynamic Neuromorphic Asynchronous Processor (DYNAP-SE) [7], or the BrainDrop chip [5].

In these systems input signals are typically represented as trains of digital pulses. These spikes are integrated by the synapses and converted into currents. The outputs of multiple synapses are then summed together and, in many implementations, are integrated by current-mode log-domain filters. The resulting current, which represents a weighted sum of the inputs, is then fed into the neuron circuit, which integrates it and produces a spike, if the integrated signal exceeds the neuron’s spiking threshold. In most cases all these circuits are passive (i.e., there is no active -always on- component), and if there is no input data, there is no dynamic power consumption. For this reason, this approach is particularly attractive in the case of applications in which the signals have sparse activity in space and time.

### 2.1 Mixed-signal analog/digital processors

The brain-inspired computational strategies proposed in this paper apply to all types of heterogeneous neuromorphic processing systems. However, here we demonstrate their benefits using the Scalable Dynamic Neuromorphic Asynchronous Processor (DYNAP-SE) neuromorphic SNN chip. This neuromorphic processor, originally proposed and fully characterized in [7], comprises both analog subthreshold circuit to emulate the neural and synaptic dynamics, asynchronous digital circuits to route spikes among the neurons and program different network connectivity schemes, and local CMOS memory cells distributed within and between neuron elements to store network parameters and connectivity routing tables (see Fig. 1). The DYNAP-SE is a multi-core architecture, with 4 cores of 256 silicon neurons each, and an asynchronous inter-core and inter-chip hierarchical routing scheme. The input and output spikes are encoded as address events and are transmitted across cores using the Address-Event Representation (AER) communication protocol [55, 56]. The silicon neuron circuits reproduce the dynamics of the Adaptive-Exponential Integrate and Fire (AdExp-I&F) neuron model [29, 57], and the synapses use Differential Pair Integrator (DPI) current-mode circuits [29] to reproduce synaptic dynamics with tunable time constants that range from micro-seconds to hundreds of milli-seconds. Each neuron has a local Content Addressable Memory (CAM) block, containing 64 12 bit addresses representing the identities of the pre-synaptic neurons that it is subscribed to. Input address-events are extended by local pulse extender circuits and converted into weighted currents that are then summed into the DPI synaptic dynamics blocs. Four different synapse types can be chosen for each neuron: AMPA (fast excitatory), NMDA (slow, voltage-gated excitatory), GABA-b (inhibitory, subtractive), and GABA-a (inhibitory, shunting). Each synapse type is implemented with a dedicated DPI circuit and independent parameters.

**Figure 1:**
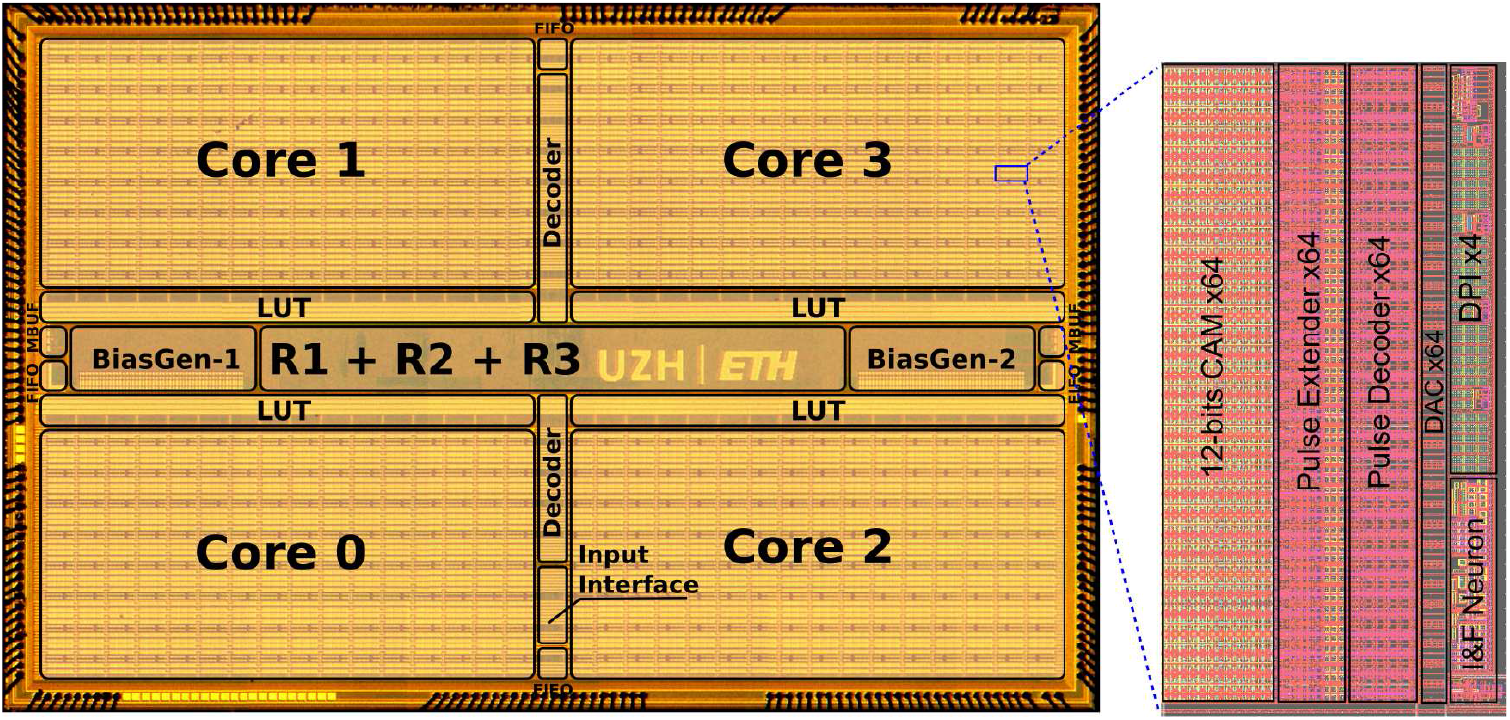
Die photo of the DYNAP-SE multi-core neuromorphic processor, with single neuron element highlighted. The chip was fabricated using a standard 0.18 *μm* 1P6M CMOS technology. Neurons between cores and between different chips can be interconnected by programming the on-chip CAM memory cells and the asynchronous digital routers labeled R1, R2, and R3. Analog parameters can be set independently for each core using on-chip 12 bit analog-to-digital bias generators.

The synapse and neuron parameters are programmable via a 12 bit temperature-compensated on-chip bias generator [58]. All parameters are globally shared within a core, and there are four independent bias generators per chip (one per core). Due to the analog nature of the silicon neurons and their device mismatch, the shared parameters used to set temporal and functional characteristics of the circuits, such as refractory period, synaptic efficacy, and time constants, vary between individual neurons and synapses. For this work we used a board with four DYNAP-SE chips, a custom Field Programmable Gate Array (FPGA) device, and a USB interface to a standard PC for configuring circuit parameters, setting up synaptic connections, sending input events and reading out neural firing activity. The board allows real-time measurement of the spiking activity of all the 4096 neurons in the board, measured as address-events, via the AER protocol, and of the analog neuron membrane potential of one (user addressable) neuron per core, with a total of 16 parallel voltage traces.

### 2.2 Device mismatch effects in neuromorphic circuits

Variations in silicon doping or mismatched geometries is intrinsic to the fabrication process of CMOS devices [31]. Device mismatch yields heterogeneous electrical properties that affect the behavior of the analog circuits, even if they have identical geometries at design time. Figure 2 shows measurements characterizing the device mismatch effects on the response properties of the analog neuron and synapse circuits implemented on the DYANP-SE. For example, when injecting the same input current to multiple instances of integrate and fire neurons of one core, the integration and spike generation circuits produce different delays in the time-to-first-spike (see Fig. 2a). Similarly, when multiple synapses that share the same weight parameters are stimulated, the device mismatch in these circuits affects the height of their response (see Fig. 2b). When measured across a multiple instances of synapse and neuron circuits belonging to the same chip, the response properties of the neuromorphic circuits that we design produce distributions which have typical coefficient of variations (CVs) ranging from 10% to 20% (e.g., see Fig. 2c and Fig. 2d).

**Figure 2:**
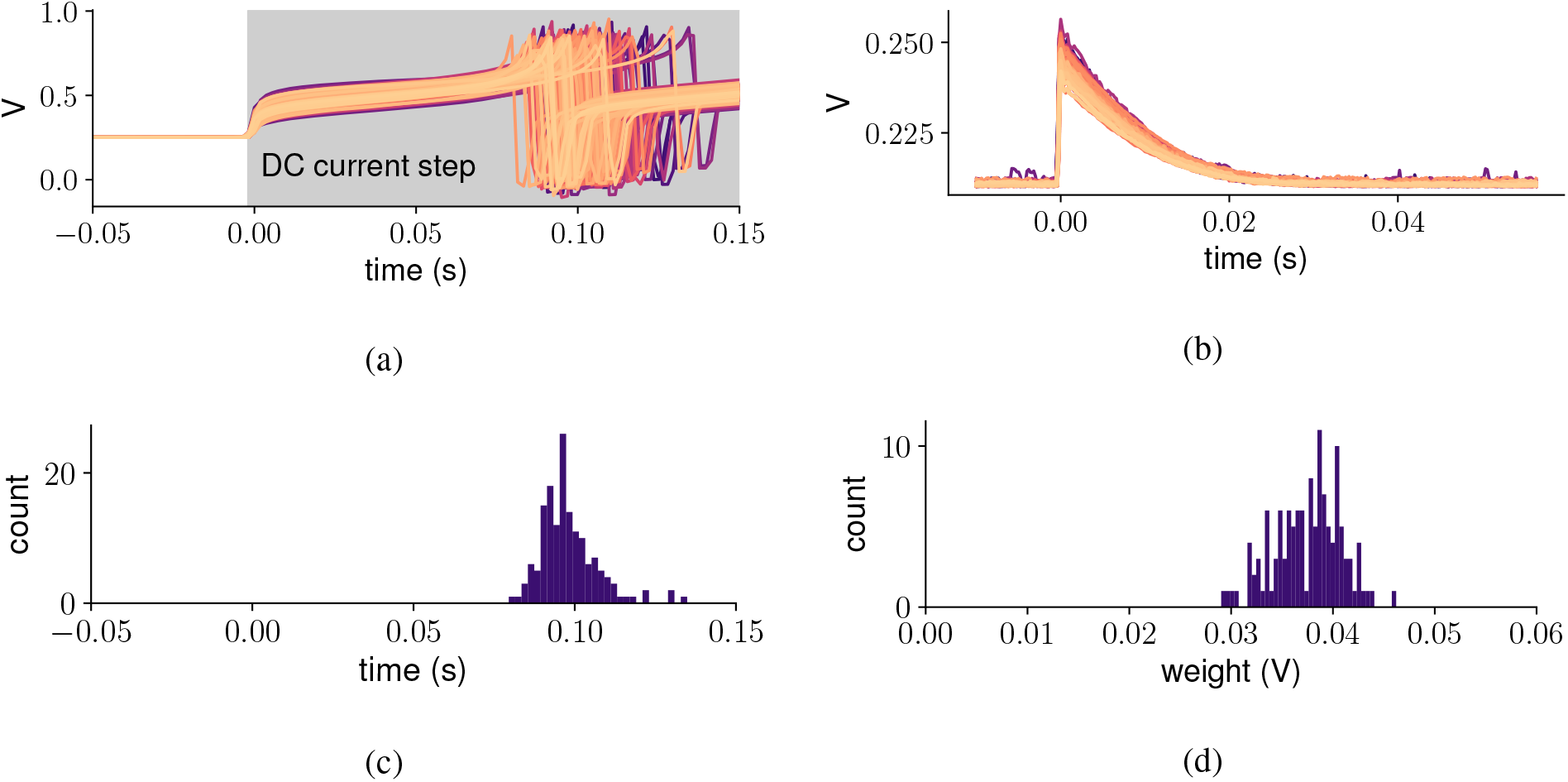
Neuron and synapse variability. (a) Aligned membrane voltage recordings of 256 neuron circuits of one DYNAP-SE core, sharing the same parameters, in response to a common DC step input current. Due to device mismatch, the time-to-first-spike of each neuron is distributed normally with CV=9%; (b) Aligned impulse responses of 256 synapse circuits of the same core. Due to device mismatch the height of the impulse response is distributed around a mean value that is proportional to the same synaptic weight parameter; (c) and (d) show the distributions of the measurements. The variation of responses is the cumulative result of individual properties distributions.

To quantify the device mismatch effects in the DYNAP-SE circuits accurately, we first defined a set of shared parameters that produce the desired average neuron and synapse behaviors and then systematically measured the circuit responses across all synapse and neurons circuits integrated on the chip. We automated the data acquisition process using a computer-controlled oscilloscope, and measured the analog subthreshold membrane response of the neuron, in response to different types of inputs. We recorded all the measurements on a work-station and carried out standard signal analysis routines to derive the neuron refractory period, the neuron time constant, the synaptic weight and the synaptic time constant from the recordings (see Fig. 3).

**Figure 3:**
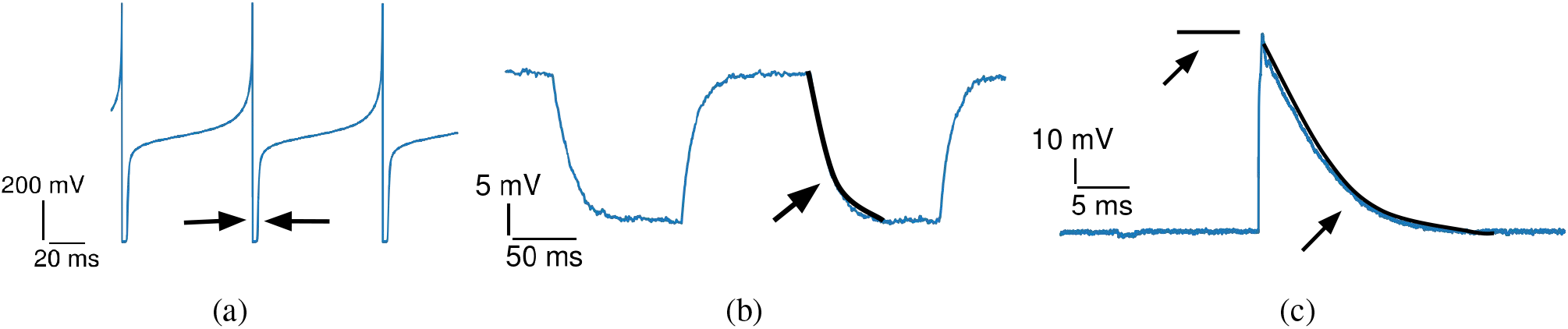
Automated measurement of neuron and synapse silicon circuit properties. (a): Measurement of refractory period of a neuron responding to constant input DC current. (b): Measurement of neuron membrane time constant by fitting the relaxation part of the neuron response after the subthreshold DC current input. (c): Measurement of amplitude (weight) and time constant of an excitatory synapse.

The refractory period of individual neurons was measured by driving the neuron to produce regular spikes trains with constant input currents, as depicted in Fig. 3a. The refractory period was defined as the time interval between the action potential reset and the voltage of the neuron rising above the 20% of its value at rest.

The neuron’s time constant was estimated by fitting its response to a step input current small enough to keep the neuron’s membrane potential below its firing threshold. The fit with was performed for the decaying part of the circuit response, corresponding to the removal of the input current (see Fig. 3b).

The synaptic weight and time constants of the synapse were estimated indirectly, by analyzing the neuron membrane potential, using a more elaborate protocol: we first set the neuron time constant to very short values, to allow the neuron to follow faithfully its input currents; then, to generate excitatory or inhibitory post-synaptic potentials (EPSPs and IPSPs, respectively) we stimulated the neurons with 20 Hz spike trains and analyzed the impulse response decay in-between input spikes (see Fig. 3c). The weight values were estimated by measuring the PSP amplitude at the onset of the input spike, and the time constant was estimated by fitting the curve with a decaying exponential.

According to the measurements performed, the neuron time constant parameter has a CV of 18%, and it’s refractory period of 8%. Similarly, the CV of the synapse time constant ranges between 7% and 10% depending on its type (NMDA or AMPA) and the weight parameter from 14% to 30%. The detailed distributions of these parameters measured across 256 neurons of one core are shown in Fig. 4. The figure illustrates how neuron refractory period, neuron membrane time constant and excitatory synapse weight distributions change for four different parameters settings, controlled by changing the values of their respective biases currents. As shown, the CV of such distributions tend to stay constant, across different parameter settings. This allows us to determine exact mismatch measures for individual properties across every core.

**Figure 4:**
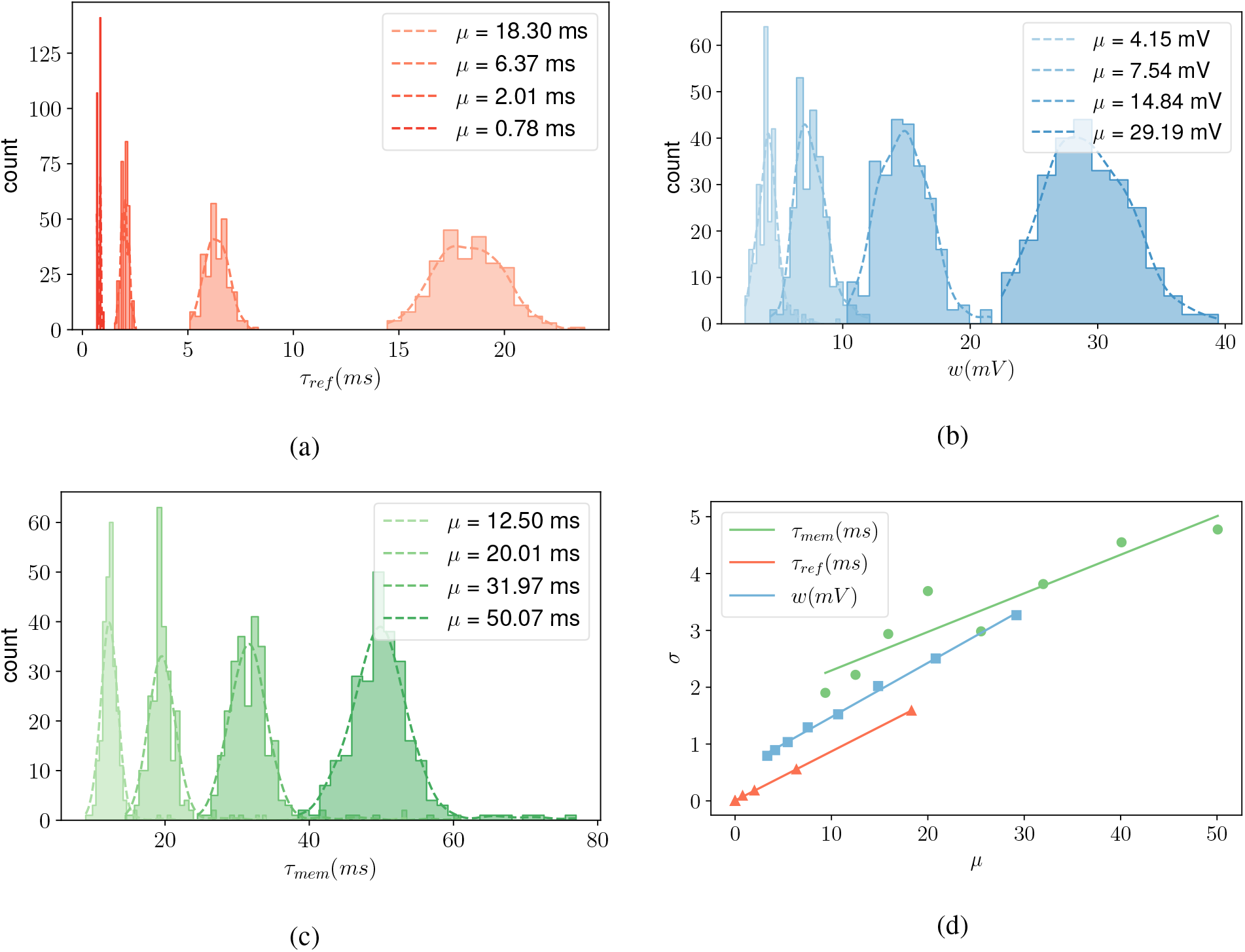
Variance behaviour across different properties of the DPI neuron circuit: (a) neuron refractory period, (b) synapse weight and (c) neuron time constant measured across 256 neurons of the same core for 4 different bias values (shown in shades of the respective color). The plot in (d) shows how the variance changes with the mean, confirming that the CV remains approximately constant for all these parameters.

Even though the variability of different parameters can exhibit different spatial distributions across the chip area, as evidenced by the patterns shown in Fig. 5, these spatial distributions differ for each parameter, and are generally uncorrelated between parameters. As a consequence, the superposition of these effects results in a heterogeneous neuron firing behavior that has no particular correlation with the location of the neuron on the chip layout, or with the specific chip used.

**Figure 5:**
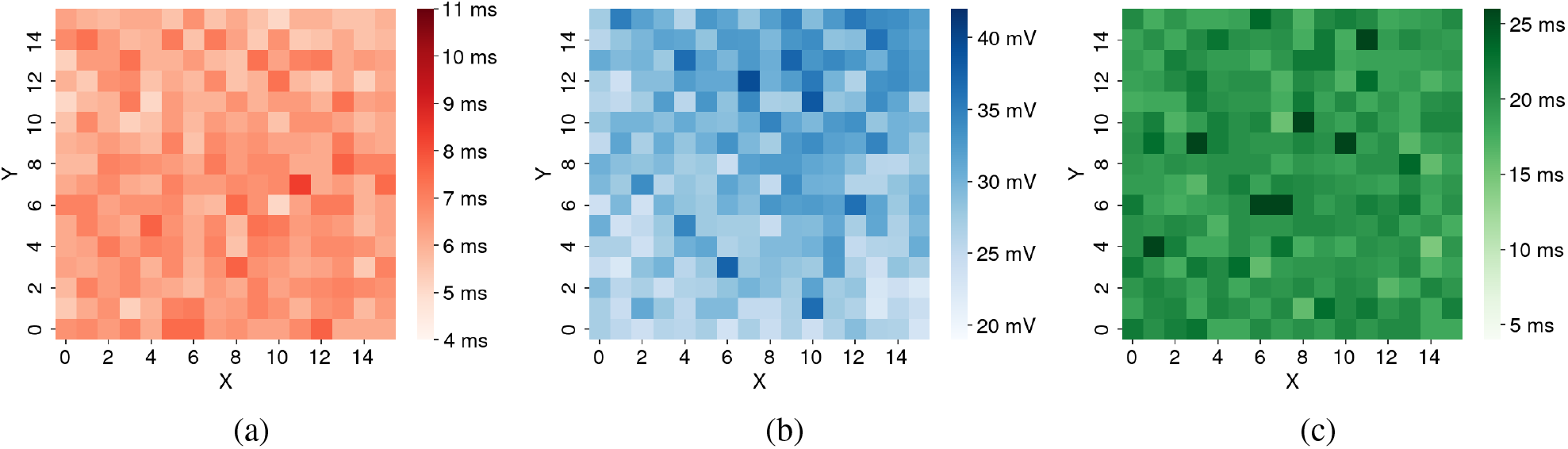
Spatial distribution of mismatch profiles for the neuron refractory period (a), the synaptic weight (b), and the neuron time constant (c). The *X* and *Y* axes represent the neuron id across the layout of the measured core.

Figure 6 shows the combined effect of both synapse and neuron variability: the plot shows the average firing rate of 16 neurons belonging to the same core, and sharing the same parameter settings. Each neuron is stimulated via its own excitatory synapse, driven by the same set of input spike sequences of increasing frequency. Although the variability of both synapse and neuron parameters produces response profiles with large differences, the ensemble average response follows the desired profile with good linearity.

**Figure 6:**
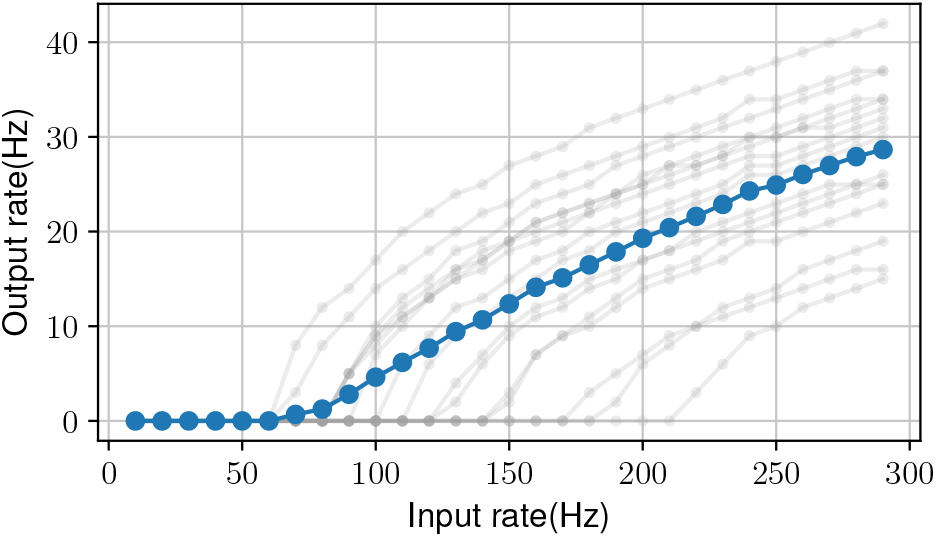
Firing rate of 16 neurons, belonging to the same core thus sharing the same parameter settings. All neurons are driven by the same series of regular spike trains with constant rates over one second and step increments of 10 Hz.

## 3 Neural processing strategies for robust computation

All physical systems that represent variables with analog signals (including electronic and biological ones) are susceptible to the effects of noise and variability. An effective strategy that can be used to improve their accuracy, is to increase the signal-to-noise ratio by resorting to signal-restoration processes at every signal processing stage. In digital systems this is achieved by discretizing the signal levels to binary values and to restoring them to these discrete levels using logic gates. Many classical analog electronic signal processing systems also perform signal restoration by using high gain amplifiers and analog filters at multiple stages of the communication pipeline. This strategy however comes at a great cost of high power consumption and large area overhead.

As in electronic chip design, animal brains have evolved to minimize both metabolic and wiring costs (i.e., power consumption and area) [26, 59]. The strategies used to achieve this include the miniaturization of components, the representation of information with the use of *ensemble methods* and *energy-efficient codes* [26, 60]. The most basic ensemble method is averaging, but cortical circuits use also more elaborate neural ensemble methods which make use of population codes and canonical microcircuit neural primitives [61–63]. Importantly, the brain also exploits also plasticity and adaptation at multiple spatial and temporal scales to compensate for noise and variability. Here we will show how some of these strategies can be adopted in neuromorphic systems design to mitigate the effects of variability, or even to exploit it, for achieving robust and efficient computation.

### 3.1 Averaging across space and time

Spiking neural electronic systems made of high precision components or ideal neurons in computer simulations represent signals with high fidelity. In analog circuits very high levels of precision for representing signals can be achieved only at great costs. Alternatively, a practical solution, extensively exploited also in the brain, is to relax the high precision constraint for single computational units, the neuron, and to distribute computation in space and time. Feeding signals to populations composed of heterogeneous units with uncorrelated mismatch in their properties, as in Fig. 6, naturally reduces the effect of noise and variability in the single units. Averaging signals over time filters out irregularities in the spike sequences generated by both the input sensors, and by the neural processing parts of the system.

The first strategy, of *space averaging encoding* can be implemented, for example, by feeding the output spike train of one unit to multiple neurons in a cluster. We tested this strategy by carrying out an experiment in which a node producing a regular spike train of 200 Hz drives a population of 256 neurons, in one DYNAP-SE core, in a way to produce an average output rate of 50 Hz. We computed their individual output firing rates, during one second of recording, and used its distribution across the cluster to compute the rate CV of different size clusters, ranging from two clusters of 128 neurons, to 4 clusters of 64 neurons each, to 8 of 32 neurons, and so on, up to 256 clusters each made of one neuron. For each combination, the CV of the rate is computed from the mean and variance of the firing rate distribution across clusters, grouping neurons selected within the same single core. To measure average values of the CV we computed the mean CV over 10 reconstructions of the same configuration with shuffled (regrouped) neurons across clusters. Figure 7a shows the firing rate CV computed in this way, as a function of cluster size. Our data are consistent with the averaging theory, which states that the standard deviation and the coefficient of variation of a distribution is proportional to the 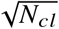, where *N_cl_* is the cluster size (see Fig. 7a). Thus, the balance between employed resources and acceptable cancellation of mismatch, i.e., firing rate CV reduction, results from a trade-off that scales with 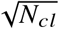.

**Figure 7:**
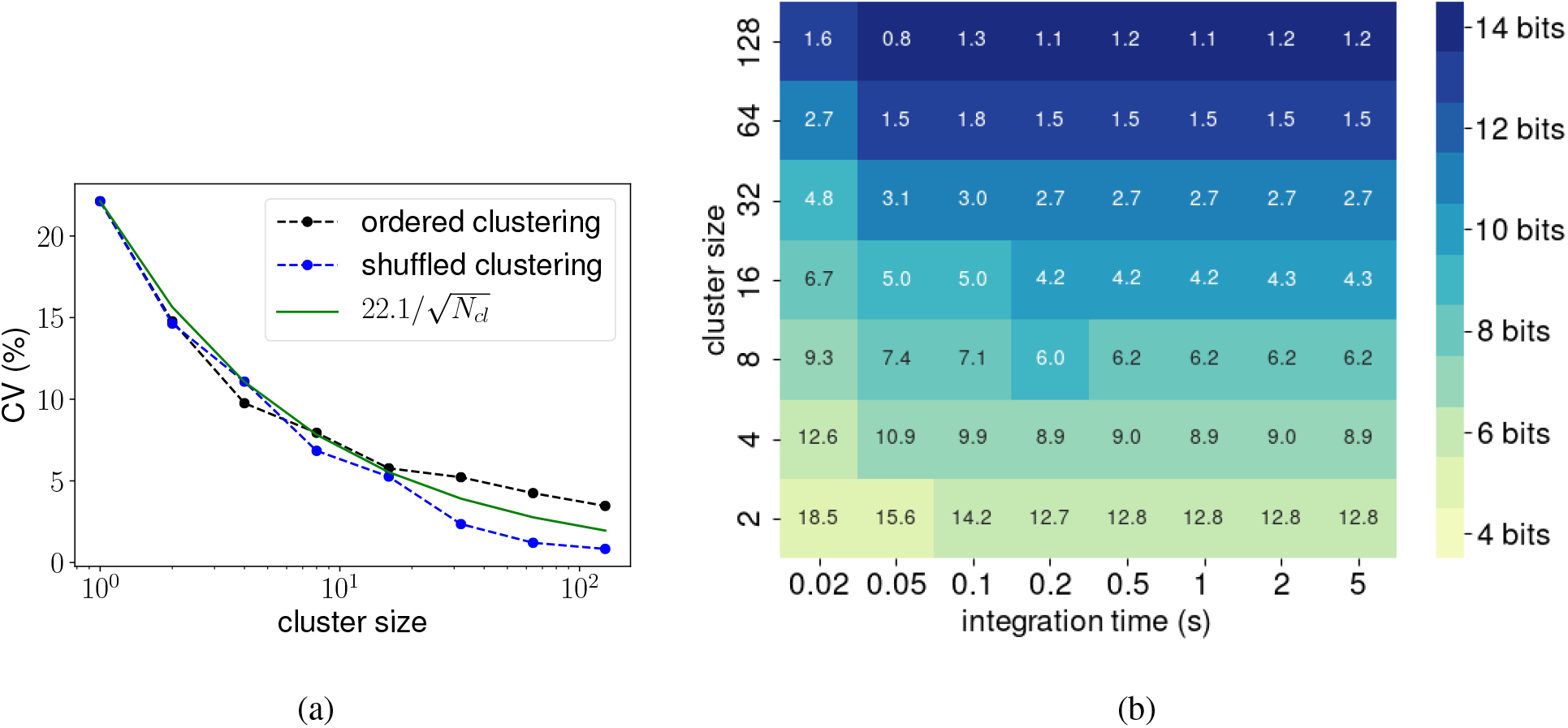
Variance of the mean firing rates of clusters of neurons decreases with cluster size. (a): recordings from 256 neurons of the chip with different cluster arrangement (solid black line for sequential and blue line for randomly shuffled clustering). All 256 neurons were used at all times, meaning the largest CV is for 256 clusters of size 1, and the lowest is for 2 clusters of size 128. The green line is a 1/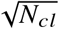 fit normalized to the first value. The integration time is 1 sec. (b) Dependence of cluster rate CV (shown in each color coded square) on cluster size (y-axis) and spike integration time (x-axis). The equivalent bit precision is for each CV value is color coded. Each CV value shown is a mean across 10 random splits (shuffles) into clusters. Darker color stands for higher precision of encoding.

In the second *time averaging encoding* strategy, the integration time window is a relevant parameter. Unlike neurons simulated with digital hardware, but very much like biological ones, mixed signal analog/digital silicon neurons produce irregular spike trains even if stimulated with constant inputs, and exhibit heterogeneity in their firing rates across multiple trials even if they input stimulus is always the same. If we fix an integration bin size, within the same bin, fast firing neurons provide more information than slow ones. In Fig. 7b, we estimate the CV for increasing bin sizes, ranging from 20 ms up to 5 seconds, calculated for each choice of cluster size. The trade-off between readout time and amount of resources is now evident: precise rate encoding requires large clusters. For example, in this experiment an equivalent bit precision of 14 bits is achieved only by using clusters that comprise 128 neurons. However, by combining longer integration times (e.g., 100 ms–200 ms) with cluster sizes of 8 or 16 neurons it is possible to achieve equivalent bit precision of 8 bits, which appears to be adequate for many artificial intelligence and neural processing tasks [64–66].

An advantage of this approach is that the chip designer does not have to make critical decisions (such as choosing the number of bits to use in a digital bus) at chip design time. The size of the cluster and the integration time can be flexibly changed and even adapted dynamically at run time.

### 3.2 Using population codes

*Encoding* signals in a robust and reliable manner is a fundamental step for processing and computing. In the nervous system this is done seamlessly at the site of sensory input and further elaborated at relay stations along the path to the central nervous system, via populations of neurons. However, to carry out robust neural computation, it is also important to choose the proper way to *represent* signals. A common strategy adopted by the nervous system is to use distributed representations [67], such as population codes [37]. Population codes are used across the nervous system to represent many types of signals, such as visual [68].or auditory cues and their spatial localization [69]. Population codes, and more generally distributed representations, are tolerant to damage, noise, and in the case of neuromorphic implementations, to device mismatch. Indeed, heterogeneity and neuronal diversity in population coding can greatly enhance the network’s information capacity [70].

A basic distributed representation can be implemented by using populations of neurons subdivided into independent clusters that represent the value of their input signal. To assess the benefits of this representation in mixed-signal neuromorphic systems we performed an experiment in which we encoded the activity of input nodes with populations of silicon neurons arranged as uncoupled clusters (see Fig. 8a). Each cluster comprises a set of neurons that are not interconnected, but that receive the same input spike train. Figure 8b shows the response of the neurons to a spatially distributed bump signal. As the neurons are driven with strong synaptic input weights, the transfer function of each neuron is close to identity for low input rates, but it saturates to a maximum rate of about 40 Hz (fixed by the neuron’s refractory period parameter of Fig. 3a) for higher input rates. This results in a suboptimal reconstruction of the bell shaped input pattern, that can be resolved using more elaborate network connectivity schemes (as we will see in the next sections). Forcing the silicon neurons to saturate at relatively low firing rates is useful for limiting the power consumption of the system. However this limits the neuron dynamic range and their ability to encode input signals that have a high dynamic range.

**Figure 8:**
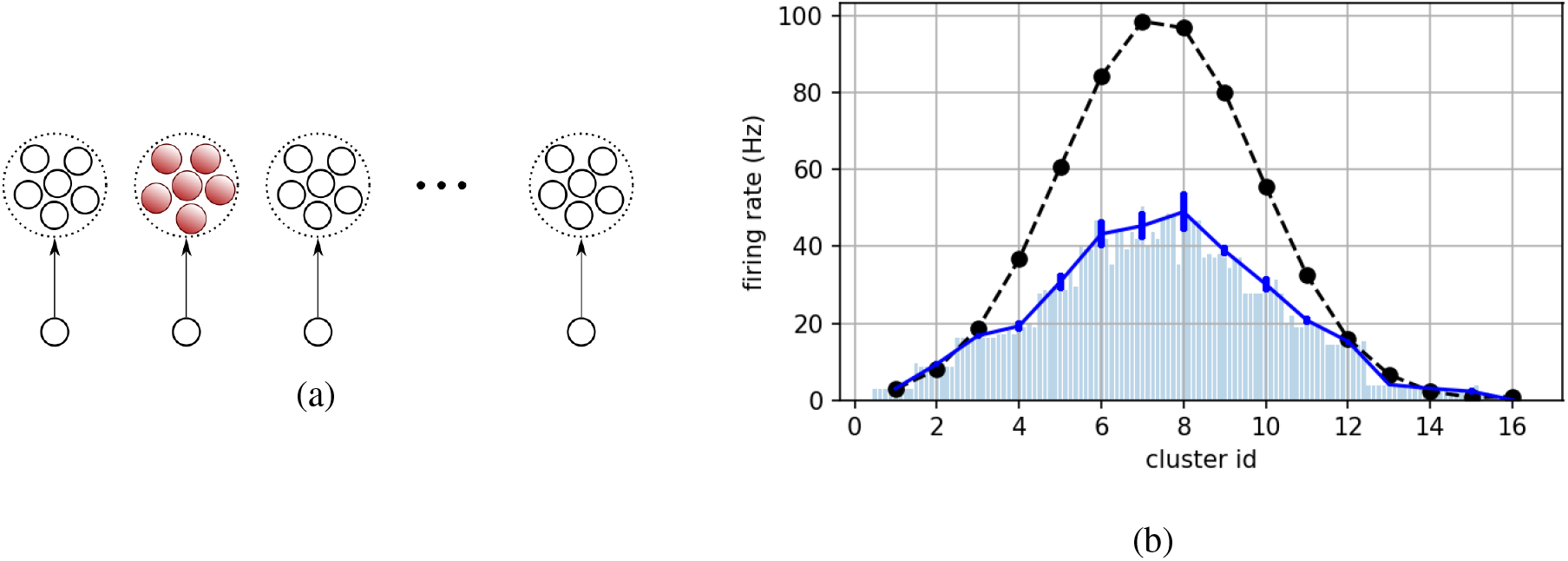
Multiple decoupled clusters of neurons representing a distributed population code. (a) Network diagram. Ten independent clusters, of 8 silicon neurons each, receive spikes from individual input nodes whose firing rates change smoothly with spatial position. Each neuron in a cluster receives the same spike sequence and contributes to the cluster’s average firing r ate. (b) Mean firing rate of the input nodes (black) and average output population activity (dark blue). Error bars indicate the standard deviation within each cluster. The light blue bars represent the firing rate of individual neurons.

This is a problem that also biological neurons face: real neurons have low firing rates and limited dynamic range compared to the signals they must represent, especially in the early sensory stages. However, using populations of heterogeneous neurons to average out the effects of variability can solve this limited dynamic range problem: as postulated in [71], populations of *N* integrate and fire neurons can faithfully encode band-limited signals that have *N* times the bandwidth of individual neurons. So resorting to the use of populations of neurons for representing signals while reducing the effect of variability has the added benefit of allowing the system to represent signals with high dynamic ranges that exceed the range of individual neurons. Furthermore, an important requirement of this theory is that neurons do not start from identical initial conditions [71]. So using populations of *heterogeneous* neurons, such as those implemented with analog circuits and/or with memristive devices, naturally satisfies the requirements of the theory. To validate this theory with our neuromorphic circuits we carried out an experiment in which we stimulated with a single Poisson input spike train a population of silicon neurons in a cluster of 16 units. The input node was configured to spike with an average firing rate of 100 Hz, while all neurons in the cluster produced firing rates with a maximum value of about 40 Hz. Figure 9 presents the experimental measurements. As shown in panels 9a–e, although individual neurons in the cluster cannot reproduce the fine details of the high dynamic range input signal, the population average (represented by the red line in the raster plots) can follow the input reliably. There is strong evidence that also real neural systems use this strategy, for example to encode signals in the vestibular system [72].

**Figure 9:**
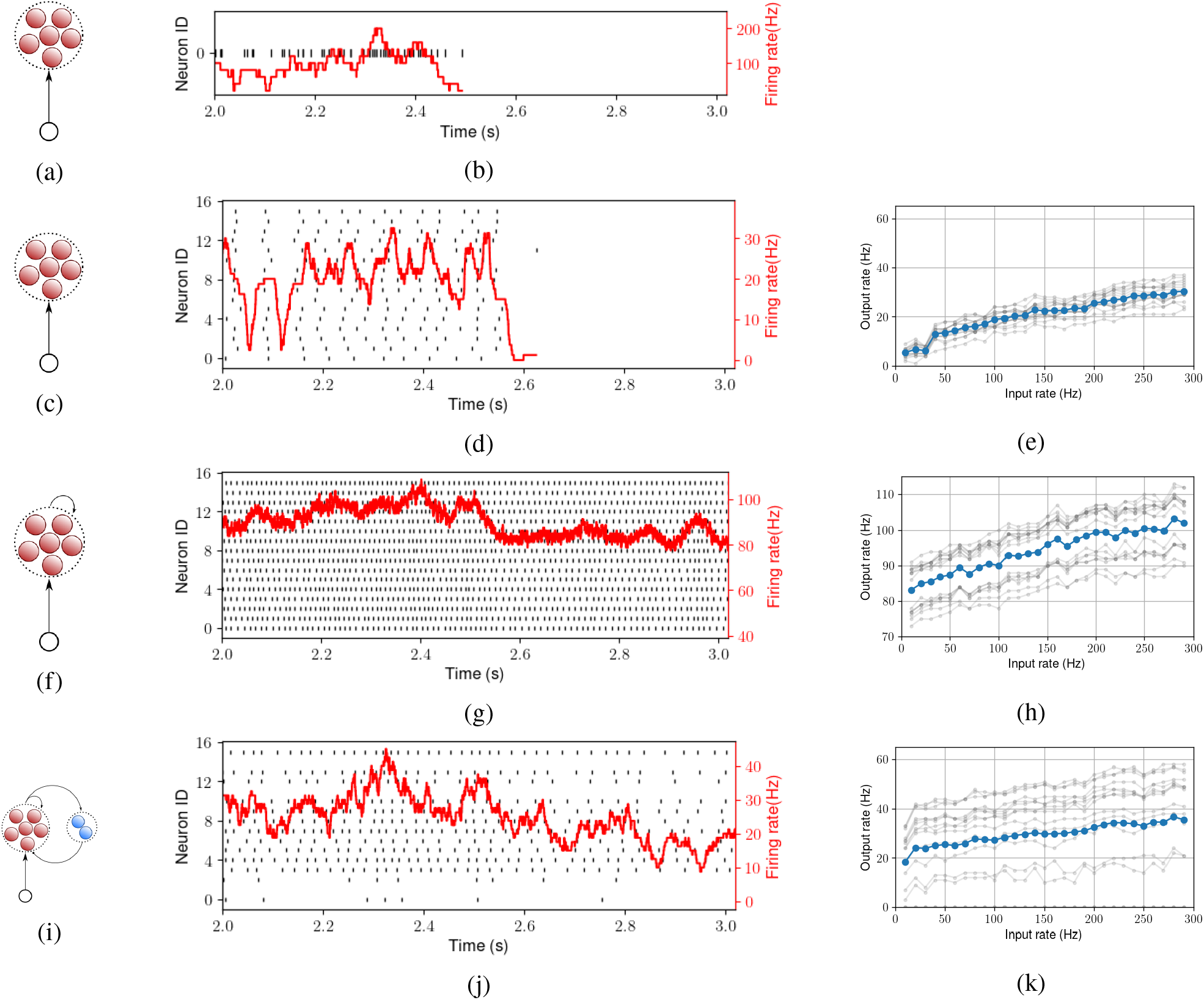
Spiking response of a cluster of 16 neurons to the same Poisson input spike train (panel b) in 3 different connectivity configurations: pure feedforward (c), with recurrent excitation (f), with additional inhibitory feedback (i). Panels on the right (e,h,k) show the response firing rates of individual neurons (gray traces) and the mean cluster firing rate (blue line) when feeding them with a regular spike train at different firing frequencies. The input spike train (b) is fed to the cluster for each trial and the response rasters (d,g,j) show the spiking response both during the input and after the input ends. The mean firing rate of the cluster is shown in red.

Also in this case, in addition to averaging and reducing the effect of variability, resorting to using populations of neurons produced extra important advantages for encoding high bandwidth signals with low-bandwidth (and low power) silicon neurons.

### 3.3 Using recurrence and self-excitation

In addition to choosing the right representation, computation heavily relies on using *gain* in the processing pathway. One way to implement and modulate gain in networks of neurons is to change the strength of the synaptic weights from layer to layer. However this process, typically achieved via synaptic plasticity, can be very slow and does not support modes of operation that require fast gain changes to carry out the desired computations. Alternatively, a strategy commonly found in the nervous systems that overcomes this problem is the use recurrence, with both positive and negative feedback. By adding recurrent excitation to the population of neurons it is possible to control and modulate the gain of the network quickly, following the fast changes in the activity of neurons rather than the slow changes in synaptic efficacy of relevant synapses. Indeed, as gain modulation is directly affected by the network activity, it can happen at much higher speeds than those dictated by plasticity mechanisms.

To demonstrate how this strategy is effective also for recurrent networks of silicon neurons, we added self-excitation to the cluster of 16 neurons of Fig. 9 (see Fig 9f). This has the effect of increasing the gain of the network, driving the response of the network to much higher firing rates compared to the 40 Hz baseline of panel 9d (see red data in Fig. 9g), while being still sensitive to the changes in the input signal, as evidenced by the linear input-output relationship measured in Fig 9h. However, as this gain modulation is obtained via positive feedback, it can be difficult to control. For example, in this case and with the parameter settings chosen, the network maintains its activity in a high persistent state even after the input has been removed. This can be a desirable effect for developing *attractor networks* [73, 74], but it can also be an undesired effect in other cases. The brain-inspired strategy that can be used to keep this effect under control is to add a negative feedback loop in parallel with the positive feedback one. This is achieved by projecting the activity of the excitatory neurons to a population of inhibitory neurons that in turn inhibits back the excitatory population (see Fig. 9i, j).

### 3.4 Balancing excitation with inhibition

In the configuration of Fig. 9, a single common Poisson source of spikes (with ISI *CV* = 1) was used to stimulate all neurons in the cluster. This led to correlated firing in the cluster, especially in presence of recurrent excitation: the average ISI *CV* of the population spike trains in Fig. 9g decreased to 0.1. This impairs energy efficiency and is detrimental for signal encoding [75]. Recurrent inhibition can help to decorrelate firing activity and significantly enhance coding efficiency [76, 77]. This is true also for our silicon neuron network of Fig. 9i. The recurrent inhibitory feedback leads to an excitatory/inhibitory balance that has the effect of producing sparse and decorrelated activity: the average ISI *CV* of the data in Fig. 9j increased back from 0.1 to 0.37.

We initially introduced recurrent inhibition as a negative feedback loop to better control the gain of the network and reduce the average firing rates in the cluster. As this mechanisms is effective in reducing correlations among the neurons it produces additional beneficial effects for efficient signal encoding and for memory compression [78, 79]. In addition, the asynchronous firing state produced in this way can generate an optimal noise structure, enabling the network to track input changes rapidly [80, 81].

### 3.5 Using soft Winner-Take-All networks

By combining the recurrent inhibition mechanisms used to sparsify neural activity of Fig. 9i with the distributed representation population coding scheme of Fig. 8 we can implement networks that exhibit both competitive and cooperative features, and that can support a wide range of useful computational features.

Figure 10a shows an example of such a network: cooperation is mediated by excitatory connections with local connectivity (nearby clusters are connected via excitatory connections), and competition is achieved by means of global inhibitory connections: all clusters are inhibited by the population of inhibitory neurons, which are stimulated by the excitatory neurons of all the clusters in the network. These types of networks are often referred to as soft Winner-Take-All (sWTA) networks. In these networks nearby neurons are biased to have similar responses properties (e.g., similar stimulus preferences, or receptive fields), and thus create a map in which close by units represent similar features.

**Figure 10:**
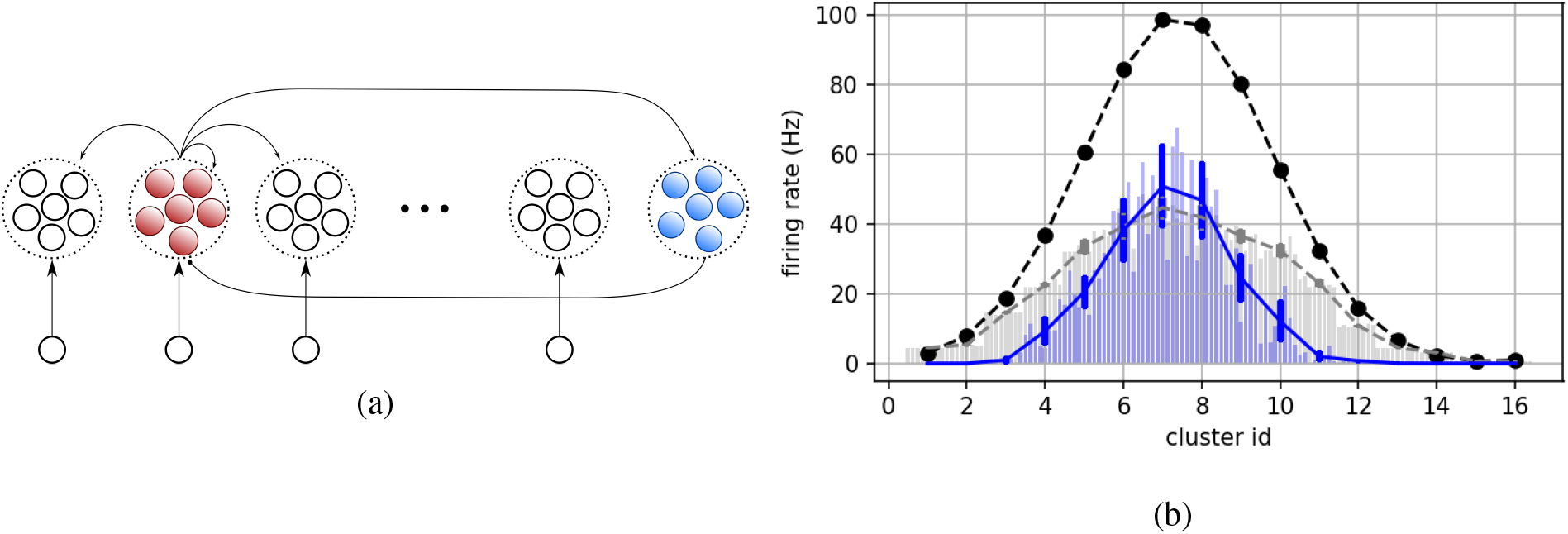
Example of a competitive network comprising multiple clusters coupled via local excitation and global inhibition; (a) network diagram of the network which comprises 16 clusters of 8 excitatory neurons each and a global inhibitory cluster of 20 neurons. Input nodes stimulate all neurons in the cluster with equal strength; neurons in each cluster have all to all recurrent connections (not shown in the diagram) with 100% connection probability; neurons of neighboring clusters are connected with a 10% probability; excitatory to inhibitory neurons have a 20% connection probability and inhibitory to excitatory neurons have an 80% connection probability. (b) network activity measured experimentally from the chip in response to a Gaussian input bump of Poisson spike trains with mean amplitude of 100Hz (black dashed line). The gray dashed line represents the response of a pure feed-forward network without recurrent excitatory and inhibitory connections. The solid blue line represents the competitive network response, with recurrent excitation and global inhibition activated. The error bars in this data reflect the variability within c lusters. The vertical bars of both gray and blue plots represent the responses of the individual neurons in the cluster.

Depending on their parameters, the same sWTA network can be used to process continuous signals and represent different features that change smoothly in feature space (e.g., the orientation of a visual stimulus), or to manipulate discrete symbols such as numbers and numerable variables (e.g., one of n possible keywords). The recurrent connections in the network make the outputs of individual clusters depend on the activity of the whole network, and not just on the neurons driven by the local input [82]. As a result, sWTAs can perform both linear operations, such as amplification by a linear gain or locus invariance, and complex non-linear operations, such as normalization, selective amplification and non-linear selection, multi-stability, or signal restoration [42]. Interestingly, it has been observed that, despite significant variation across cortical areas, sWTA types of connectivity patterns are found throughout all of the neocortex [83, 84]. Indeed, this architecture is a “canonical microcircuit” that can be used as a fundamental computational primitive for multiple types of both signal processing and computing tasks [42]. It has been shown that the computational abilities of sWTAs are of great importance in tasks involving feature-extraction, signal restoration and pattern classification problems [41]. Artificial neural networks with this architecture, and their neuromorphic hardware implementations, have been used to detect elementary image features (e.g., oriented bars) and reproduce orientation tuning curves of visual cortical cells [85–87].

#### Tuning curve sharpening

Figure 10b shows the experimental measurements from the sWTA network formed by the silicon neuron circuits of the DYNAP-SE chip, with (blue solid line) and without (gray, dashed line) the recurrent connections. The black dashed lines in Fig. 10b represent the input to the network. As for Fig. 8b the output firing rate of the silicon neurons is kept low via the refractory period setting, to reduce power consumption. So while low frequency inputs are followed faithfully, high frequency ones are scaled to lower values. However, what is important is the effect of the recurrent inhibition: this network of silicon neurons can reproduce the selective amplification features expected from sWTAs models, amplifying with a gain higher than one the strongest inputs while at the same time suppressing the weaker ones. This gives rise to a “sharpening” of the tuning curve similar to what has been measured in real cortical cells [82, 85, 88].

#### Selective amplification

Due to their cooperative/competitive nature, clusters with the highest response are amplified while weaker ones are suppressed. To highlight the selective amplification properties of the sWTA presented in Fig. 10 we stimulated the network with two input bumps of slightly different amplitudes. Figure 11 shows the response of the network to these inputs. The reference response of the feed-forward network without the recurrent feedback is shown in Fig. 11a. As expected, the network follows the input with two bumps of the same width and proportional amplitudes. As soon as feedback is enabled, the non-linear processing features of the network become evident: higher inputs are preserved and “pass through” to further processing stages, while (even slightly) weaker inputs are almost fully suppressed (see panels (b) and (c) of Fig. 11 for the cases in which the stronger input is on the left or right side, respectively).

**Figure 11:**
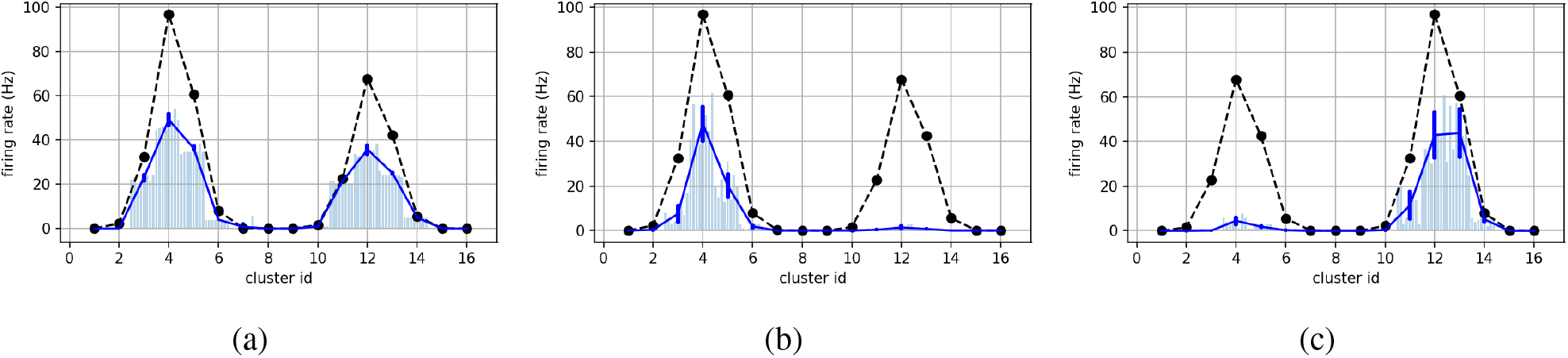
When provided an slightly different inputs (two bumps with amplitudes of 100Hz and 70Hz), the sWTA network of Fig. 10 selects the strongest one. Panel (a) shows the network’s pure feed-forward response when both recurrent excitatory and inhibitory connections disabled. The dashed line and solid circles represent the input firing rates; the solid blue lines represent the activity averaged over clusters of 8 neurons each; the histograms represent the firing rate of individual neurons. Panels (b) and (c) show the response of the network when the recurrent connections are activated: a stable output activity bump forms around the location of the stronger input, while the response to other input is almost completely suppressed.

#### Signal restoration

These very same features enable the network to support another important computational primitive: that of “signal restoration”. This is the very process that allowed the success of logic-gates in digital computing systems, always restoring their output to a nominal “1” level or “0” level. If signals are encoded with distributed population codes as provided by the output of sWTA networks, then multiple layers of such networks automatically carry out signal restoration. To demonstrate this experimentally we provided in input to the hardware sWTA network a “bump” signal as produced by neurons belonging to another sWTA network, and corrupted it in two different ways: with “dead” neurons (i.e., by silencing completely 30% of the input units) and with noisy neurons (i.e., by adding 20% noise to the nominal value of each input unit). As shown in Fig. 12 the network is able to recover the original population encoded signal and produce an output that is very close to the un-corrupted input.

**Figure 12:**
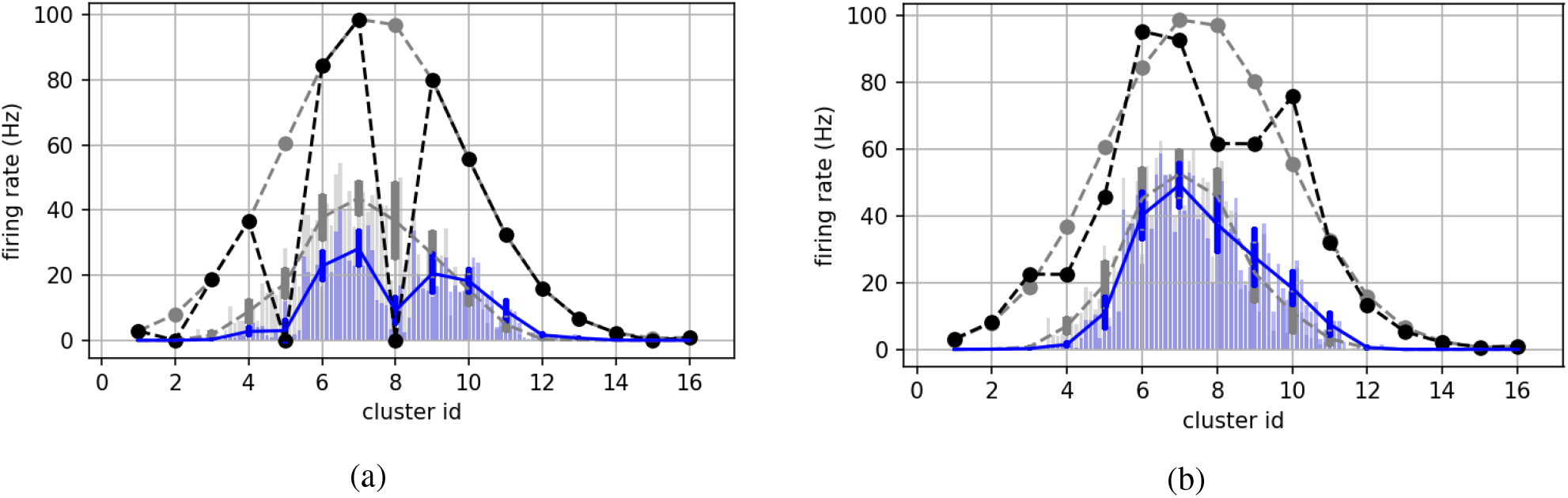
Signal restoration demonstration. The setup is identical to Fig. 10. (a) response of the network to an input bump with 30% of rate inputs missing completely. (b) response of the network to an input corrupted by 20% Gaussian noise. The solid black line shows the perturbed input overlaid over gray ideal input and response. In both cases blue curve shows the response of clusters of the network. The blue error bars indicate SD withing each cluster. The blue histogram bars show activity of individual neurons. In both cases the resulting activity bump location still represents the intended population code.

#### Attractor dynamics

sWTA networks can exhibit dynamic characteristics of attractors underlying many cognitive functions, including decision making and working memory. Depending on their connectivity patterns, sWTA networks can open boundary conditions (e.g., to represent a variable ranging from zero to a maximum value) or closed boundary conditions (e.g., representing an angle that can take values between 0 and 360°). In the latter case the sWTA network forms a ring attractor, rather than a line in the network state space. Both types of configurations have been found in different brain areas (e.g., in the brain stem for oculomotor control [89], or in the fly’s central complex for navigation [90, 91]). In both types of networks, the position of the neuron, or cluster of neurons, that has the highest activity represents the value of the variable being encoded. When driven by external signals, this activity bump stabilizes around the strongest input. Ideally, when the input is removed, and when the sWTA is configured to produce working memory behavior, the activity bump should persist and remain at the same position. However, in our experiments, after the input is removed, the bump of activity starts to drift (see Fig. 13). In these experimental results, the sWTA activity bump drifts randomly, continuously shifting between semi-stable positions across the whole population for the first two seconds of the recording. At *t* = 2 *s* an input bump is presented with a Gaussian profile. The peak of the Gaussian is indicated by the red line in Fig. 13. As evidenced, the network activity quickly shifts to the strongest input’s location and stays there until the next input is presented. After the input is removed again (at *t* = 4 *s*) the bump starts to drift again. Thus, this sWTA does not have a single, strong attractor able to immediately cancel the memory of an input. This phenomenon has been modeled also in theoretical works, and has been shown to be controllable by endowing the network with homeostatic plasticity features [92], which can be readily implemented in neuromorphic electronic circuits [93, 94]

**Figure 13:**
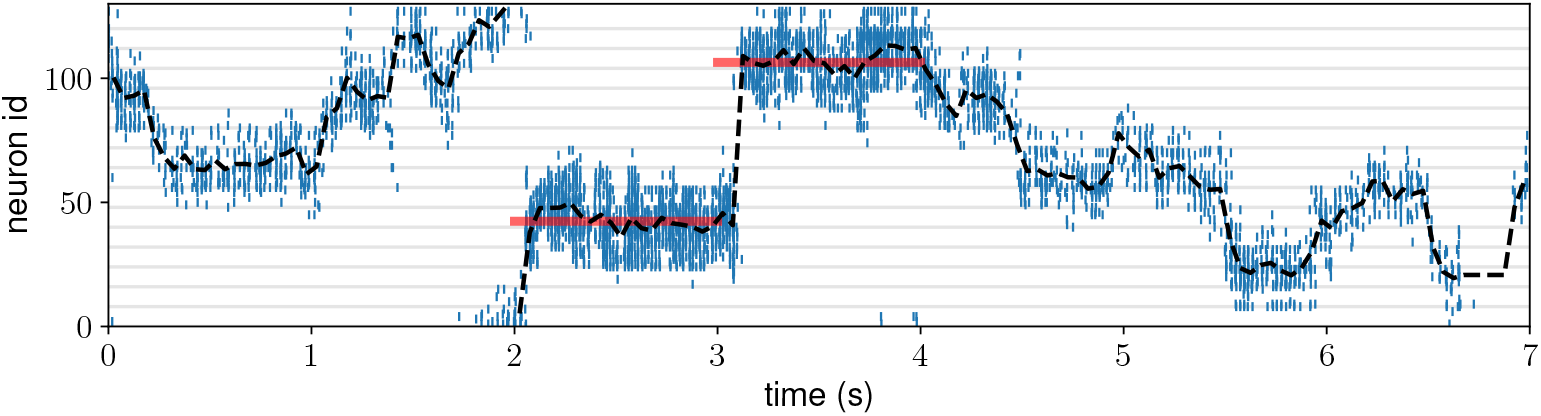
Population code follows the input and then drifts freely when no input is presented. A WTA network identical to Fig. 10 receives two population coded inputs with 1 second length (red lines). The network activity (blue spikes) is focused by input, and then begins to drift when the input is removed. Black dashed line shows the population vector (the population coded value) estimated every 20 ms. Grey horizontal lines illustrate division of neurons into clusters of 8.

### 3.6 Linking multiple soft Winner-Take-All networks together

sWTA networks have been associated to canonical microcircuit that sub-serve many computational properties in many cortical areas [95–97]. As these microcircuits have been found to be often reciprocally connected in the cortex [98], it is natural to hypothesize that multiple sWTA networks coupled among each other have the potential of performing even more complex computations.

Indeed, it has been shown how these coupled networks can form arbitrary relations between the variables encoded in the individual sWTA populations [99], or can self organize to learn relations when coupled and endowed with spike-based learning mechanisms [100]. One example of a relational network formed by coupling multiple population-coded variables among each other is the model of sensory-motor mapping between head and eye positions [101]. Another example of a relational network is shown in Fig. 15, where three sWTA networks are coupled to form the relationship *A* + *B* = *C* [99, 102]. The neuromorphic implementation of this relational network consists of 4 distinct populations of silicon neurons: three input 1D sWTA networks, representing the variables *A, B*, and *C*, and one hidden 2D population encoding the relationship between the three variables (see Fig. 14a). The links from the 1D networks to the 2D one are bi-directional, therefore creating a recurrent network that allows the activity of 1D networks to influence the others. We configured the hardware such that the variables encoded by the networks range from 0 to 1, and we implemented closed boundary conditions, such that the variables are wrapped around (i.e., if an operation increases a variable by *n* beyond 1, the result is only a fractional part of *n*). The plots of Fig. 14b shows experimental data measured from the silicon neurons representing the different networks, as populations *A* and *B* were being driven by external inputs. The choice of the relationship used to demonstrate this principle was arbitrary. Furthermore, in this example we set the parameters of the networks (and in particular the weights of the hidden population) manually, to demonstrate the proper operation of the relationship chosen. It is interesting to note though that arbitrary relationships, including non-monotonic ones, such as *A^2^* + *B^2^* = *C^2^*, can also be trained through local spike-based learning rules [100, 103].

**Figure 14:**
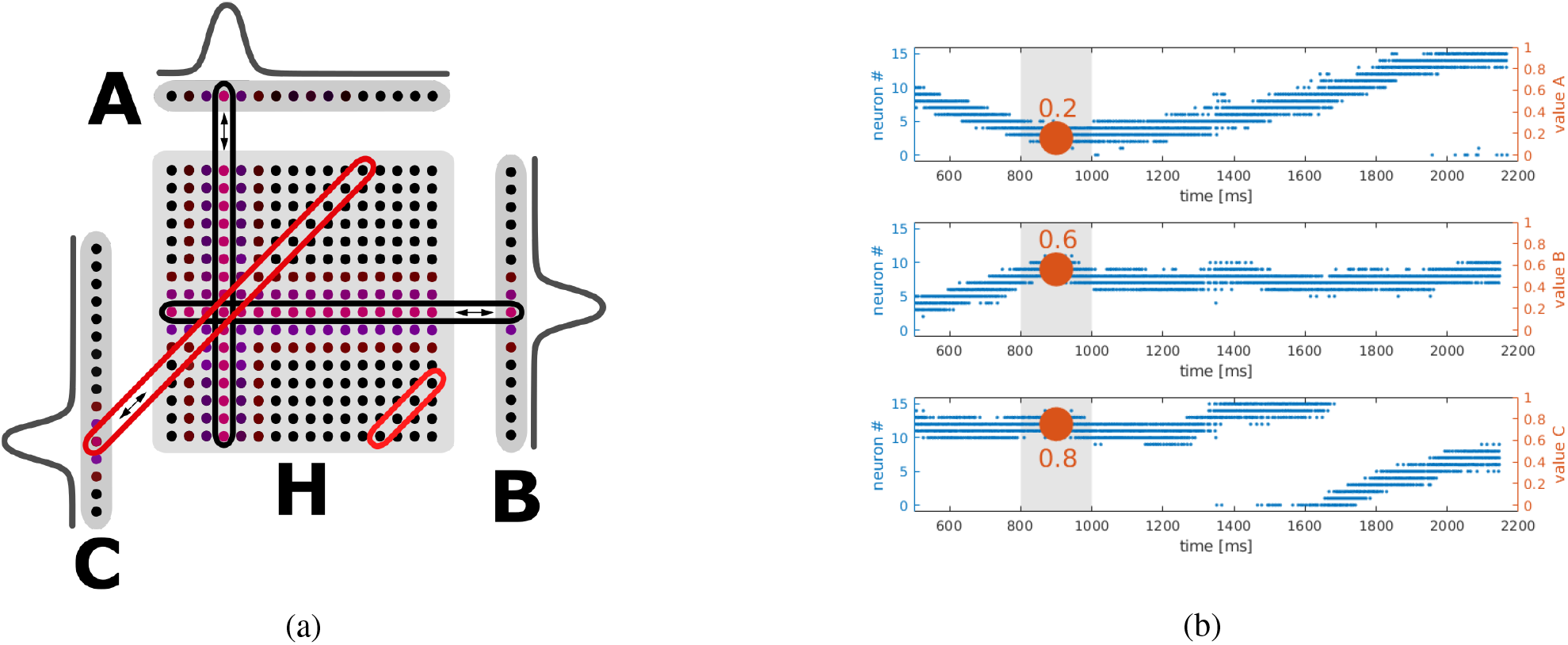
(a) Relational network implementing the relation A+B=C with three coupled 1D sWTA networks via a 2D hidden population H. (b) Raster plot of input variable populations A, B, changing over time, and output variable population C continuously holding the relation.

### 3.7 Using spike-based learning and plasticity

Perhaps the most powerful and widespread strategy used by the brain to maximize robustness for neural computation is plasticity and learning. Indeed, a large number of neuromorphic learning circuits and models have been proposed to improve robustness to device mismatch and mitigate the effects of noise in both the input signals and the system parameters [49, 54, 104–111]. Similarly, many studies on spike-based learning circuits interfaced to memristive devices have shown how plasticity and on-line learning can overcome the variability and low-resolution properties of the nanoscale memristive devices [112–115].

An increasing number of spike-based learning models compatible with neuromorphic computing technologies is being proposed, that can lead to novel memristive/CMOS robust and fault tolerant neuromorphic computing systems [116–119]. Also for these systems, variability and heterogeneity in neurons can lead to extremely powerful benefits. Booting studies have proven that, if one considers single neurons as “weak” linear classifiers, i.e., perceptrons [120] that respond to patterns they have been trained to recognize with low accuracy, then using populations of heterogeneous neurons trained to recognize the same patterns will increase the accuracy of classification and decrease the recognition error exponentially, with the population size [121].

Therefore, plasticity and learning have an extremely high potential for enabling robust computation on mixed-signal neuromorphic hardware computing substrates. However, as this is still an active area of investigation and exploring these mechanisms would go beyond the scope of this work, we refer the reader to the literature cited in the section and to a recent survey of local learning rules compatible with the constraints of neuromorphic circuits [122].

## 4 Discussion

More and more evidence is being accumulated showing that biologically plausible constraints in neural networks, such as heterogeneity [34–36], non-negativity of parameters, and partitioning neurons distinct excitatory/inhibitory cell type [123].improves robustness, reliability, and overall computing performance. Neuromorphic processing systems comprising either mixed-signal electronic circuits or memristive devices, or both, naturally support and even enforce such constraints at no additional cost (e.g., without requiring extra random number generator circuits). In this work we presented biologically inspired strategies compatible with such constraints and showed how they can support the construction of reliable artificial neural processing systems, using neuromorphic physical computing substrates that use massively parallel arrays of electronic components affected by device variability (e.g., such as subthreshold transistors, or memristive devices). Inspired by biology, we resorted to using populations of spiking neurons to reduce the effects of device to device and cycle to cycle variability. We used a mixed-signal neuromorphic processor to measure the amount of variability present in the silicon neuron circuits and characterized the trade-off of population size versus accuracy in producing reliable mean firing rate figures. Although the figures reported are specific to the particular chip used, the overall approach holds for any technology and any design that uses multiple physical instantiations of analog electronic circuits to directly implement statefull elements that emulate synapse and neural function (e.g., as in neuromorphic processing systems that use memristive cross-bar arrays) [23, 30]. Once we accepted the fact that robustness, precision, and fault-tolerance could be improved by trading off accuracy with area (as for this approach the area of the physical substrate increases with the number of neurons used in the population), we found that this strategy offered many more computational advantages than just accuracy: by using populations of neurons to represent signals, and by adopting the same strategies adopted by biological neural processing systems (namely, to use distinct classes of neurons and synapses for the excitatory and inhibitory signal pathways, to balance excitation with inhibition, and to represent signals with population codes) we demonstrated how such neuromorphic processing systems can support signal reconstruction, efficient coding, and coherency of representations. We provided chip measurements demonstrating how such population coding strategies could then be exploited to create robust and reliable computational primitives such as selective amplification and signal restoration. Additional advantages of this approach have also been demonstrated in experimental and computational neuroscience, such as ultra-fast response times of populations of real neurons [124], robust and efficient representation of time varying signals [125], and reliable propagation of information across in multi-payer networks of inhomogeneous spiking neurons [126].

If implemented with bit-precise digital circuits or simulated in software, the effects of variability and inhomogeneity and the advantages of the approaches used to cope with them could not be revealed nor exploited, unless explicitly simulated. This holds for standard computers, custom ANN accelerators, and fully digital time-multiplexed neuromorphic computing systems which integrate numerically the dynamic equations to simulate the function of multiple neurons [4, 6, 127]. On the other hand, since these digital systems support fetching data from external memory banks, they can take advantage of the high density of DRAM and implement complex large-scale systems which can accomplish extremely remarkable achievements [128], even without adopting the brain-inspired strategies and methods presented here. This however comes at the cost of exceptionally large computational resources and power consumption figures [129, 130].

The neuromorphic systems we have been discussing in this work implement “statefull neural networks” that use memory and area resources at every neuron and synapse. Therefore scaling such systems to very large sizes is still an open challenge, which might be solved in the future with the advent of memristive devices and other emerging memory technologies, but which still cannot be easily addressed today. Nonetheless, rather than replacing digital ANN hardware systems, these neuromorphic systems can complement them, for solving problems that to not require large-scale Deep Neural Network (DNN) models. Indeed, there are many practical problems that can be solved with shallow feed-forward networks and populations of perceptrons [120] or small recurrent networks with temporal dynamics and a linear read-out layer [131, 132]. For example, both the chip and principles presented here have already been applied to solving clinically relevant problems, to classify ECG and EEG signals and detect anomalies in heart beats or indicate putative epilpeptogenic zones in epileptic seizure patients [133–135]. In general these mixed-signal in-memory computing systems can provide advantages over classical ANN accelerators for those types of problems that require very limited amount of resources, in terms of volume, memory, and power, and that cannot resort to off-line cloud computing (i.e., “edge-computing” or “extreme edge-computing” applications). Indeed, the best solution will likely lie in the combination of both approaches, using the ultra low-power neuromorphic systems to carry out “always-on” sensory processing and basic classification tasks, and act as “watch-dogs” that can alert and activate the more powerful and power-hungry ANN accelerators for carrying out more sophisticated data-processing tasks with higher accuracy. The design of the optimal intelligent neural network processing system will therefore lie in the judicious combination of bottom-up, brain-inspired mixed-signal circuits, and top-down application driven digital computing system [136].

## 5 Conclusions

On our journey to achieving robust computation using inhomogeneous and noisy neural computing substrates, we found that the basic principle of averaging the activity of multiple inhomogeneous neurons led to many more computational advantages that go well beyond the expected one of reducing the variance with the square root of the number of neurons. In doing so we explored the relation between the size of the neuronal cluster size and the mean spike rate integration time, to reliably encode scalar values. We validated and quantified our studies using a mixed-signal analog/digital neuromorphic chip. Using this device as a representative example we could link the size of the cluster and the integration time to the equivalent number of bits in representing the mean spiking rate of the population. Once determined the number of neurons in a cluster required to achieve a desired precision, we added recurrent excitation and inhibition within the cluster, and configured the parameters in a way to keep neurons active also after removing the input stimulus to implement working memory. This strategy led to the reduction of correlations in the spike timings and cluster firing rates, which is a desirable feature in many models and applications. By adding local excitation and global inhibition among the clusters, we produced sWTA dynamics which produced even more interesting and robust computations, for example by reliably encoding scalar values with population coding, and by performing signal restoration. Finally we showed how recurrently coupled sWTA networks can perform non-trivial operations on these scalar values, demonstrating network-level examples neuromorphic processing systems that can be used to map sensory inputs to motor outputs, and produce state dependent computation. This work therefore demonstrates that reliable and robust computation can indeed be achieved using resource-constrained and inhomogeneous neuromorphic processing systems, by adopting the same strategies and principles used by the nervous system.

## Author contributions

G.I., S.S. and D.Z. conceived and designed the research. D.Z. developed software, performed experiments, and collected the data. D.Z., S.S. and G.I. analysed and interpreted the data and wrote the manuscript.

## Acknowledgments

This work is supported by the ERC Grant “NeuroAgents” (Grant. No. 724295) and in part by “Bando Fondazione di Sardegna 2022 e 2023 - Progetti di ricerca di base dipartimentali”.

